# Meta-gene markers predict meningioma recurrence with high accuracy

**DOI:** 10.1101/692624

**Authors:** Zsolt Zador, Alexander Landry, Benjamin Haibe-Kains, Michael D. Cusimano

## Abstract

**Background:** Meningiomas, the most common adult brain tumors, recur in up to half of cases[1–3]. This requires timely intervention and therefore accurate risk assessment of recurrence is essential. Our current practice relies heavily on histological grade and extent of surgical excision to predict meningioma recurrence. However, prediction accuracy can be as poor as 50% for low or intermediate grade tumors, which constitute more than 90% of all cases. Moreover, attempts to find molecular markers, particularly for predicting recurrence of intermediate tumors have been impeded by low[4] or heterogenous[5–8] genetic signal. We therefore sought to apply systems-biology approaches to transcriptomic data to better predict meningioma recurrence.

**Methods:** We apply gene co-expression networks to a cohort of 252 adult patients from the publicly-available genetic repository Gene Expression Omnibus. Resultant gene clusters (“modules”) were represented by the first principle component of their expression levels, and their ability to predict recurrence assessed with a logistic regression model. External validation was done using an independent merged cohort of 108 patients using the same modules. We used the bioinformatics database DAVID[9] to examine the gene ontology associations of each module.

**Results:** Using gene co-expression analysis, we were able predict tumor recurrence with high accuracy using a single module which mapped to cell cycle-related functions (AUC of 0.81 ± 0.09 in external validation). Further, the module maintained its accuracy with removal of up to 20 of it’s most individually predictive genes, suggesting robustness against noise at the single gene level.

**Conclusions:** With the easy accessibility of gene panels in healthcare diagnostics, our results offer a basis for routine molecular testing in meningioma management.

## Background

Meningiomas constitute approximately 34% of all brain tumors, affect 3% of the adult population[10] with an incidence rate of 8.36 and 3.61 per 100,000 person years in females and males respectively [11]. For over half a century, the prediction of meningioma recurrence has relied solely upon histological features[11] (World Health Organization grading from I to III) and degree of surgical excision [1]. While about 70-80% of fully excised WHO grade I meningiomas do not recur, 20-30% recur over follow-up with half of the recurrences occurring before the tenth year and half after ten years[12]. For WHO grade II lesions completely excised, histology predicts a recurrence of 50% over 5 years [2] and a disease specific survival rate of 69% over 10 years [3]. Given the inherent inaccuracy of histology to predict recurrence, all patients consequently require lifelong monitoring with expensive MRI imaging and clinical follow-up, essentially transforming meningioma into a “chronic disease”. Not only does this represent a significant financial burden but the stress and strain on the psychological well-bring of patients is substantial as they live with the constant uncertainty as to whether the tumor will recur, face future morbidity or require more treatment. It is therefore critical to develop better methods designed to assess the risk of recurrence that take into account the intrinsic biology of the patient’s meningioma.

The genetic landscape of meningiomas is the next frontier to our understanding of their biology and its relation to disease recurrence. While some studies have identified chromosomal rearrangements [13], mutations in genes TERT [14], ATK1 [15], SMO [16] and DNA methylation [8] as correlates of tumor recurrence, none of these have successfully translated into routine clinical practice. In addition, studies on transcriptomics have been limited by low signal. However alternative methodology exists to effectively capture additive biological effects and relate them to complex clinical traits [17–19]. Such methods may help extract translatable markers of meningioma recurrence.

Gene co-expression networks are an emerging technique used to detect patterns in transcriptomics by incorporating additive signal from relatively low gene expression levels [17, 18, 20]. The technique establishes gene similarity profiles based on shared connectivity and clusters into biologically similar groups called modules. A meta-gene is computed to summarize each module based on its gene expression levels and correlated to a clinical trait. Highly correlated meta-genes can then be used to predict a phenotype such as recurrence. With the affordability of gene expression profiling, meta-gene based diagnostics of tumor recurrence becomes feasible. With contribution of multiple module genes a meta-gene seeks to holistically represent a biological process. This multi-gene signature provides a robust against fluctuations of individual expression levels.

In the current study we hypothesized the existence of meta-genes consisting of gene modules that predict tumor recurrence with high accuracy. By annotating modules correlated with recurrent phenotypes we aim to provide further insight into the underlying biology driving tumor recurrence in addition to the clinical application.

## Methods

### Data collection and pre-processing

We identified 17 studies containing meningioma gene expression data in the *Gene Expression Omnibus* (GEO)[21] with four datasets (GSE43290, GSE1658, GSE74385, and GSE16181) including annotated tumor recurrence (Table 1). Gene expression analysis for these studies was carried out using microarray platforms Affymetrix Human Genome U133A, Affymetrix Human Genome U133 Plus 2.0, Illumina Human HT-12 V4.0, and SU Homo Sapiens 912, respectively. Each study was background corrected, log2 transformed and quantile normalized except GSE16181, which had previously been pre-processed. After this step, we filtered to include only genes common to all four studies. Discovery was done using dataset GSE16181 and external validation was done using GSE43290, GSE1658 and GSE74385 datasets merged with *ComBat*, a well-established Bayesian batch correction tool[22]. Notably, gene expression matrices were globally scaled to a mean of 0 and standard deviation of 1 prior to merging, as previously suggested[23].

**Table 1:** Study demographics.

| GEO entry | N patients<br>(n recurrence) | Median age<br>(95% CI) | N male<br>(%) | Median F/U<sup>†</sup><br>(95% CI) | Median TTR<sup>‡</sup><br>(95% CI) | WHO grade<br>(I II III) [n] |  |  | N genes<br>(overlap)<sup>∨</sup> |
| --- | --- | --- | --- | --- | --- | --- | --- | --- | --- |
| GSE43290 | 47 (8) | 65.0 (60.7-69.3) | 13 (27.7) | 4.7 (3.7-5.7) | 5.8 (3.6-8.0) | 33 | 12 | 2 | 12993 (12729) |
| GSE16581 | 16 (6) | 54.0 (48.6-59.4) | 7 (43.8) | 4.4 (3.1-5.7) | N/A | 8 | 7 | 1 | 20988 (12729) |
| GSE74385 | 45 (22) | N/A | N/A | N/A* | N/A | 17 | 8 | 20 | 21210 (12729) |
| GSE16181 | 252 (92) | 45.0 (43.4-46.6) | 53 (21.0) | 10.0 (9.6-10.4) | 3.0 (2.5-3.5) | 140 | 88 | 24 | 912 (520) |

<sup>†</sup> Follow-up (years)
<sup>‡</sup> Time to recurrence (years)
<sup>∨</sup> Number of genes that overlap with merged discovery cohort
\*Follow up at least 3 years for non-recurrent tumours, though specific times are not available

### Gene module detection and meta-gene computation

Our analysis was based on the well-established Weighted Gene Correlation Network Analysis (WGCNA), which has been described in detail elsewhere[18]. We first constructed an adjacency matrix with gene-gene Pearson correlations. Per convention, soft-thresholding was introduced by raising correlations to a common power (a hyperparameter, ß), which was selected as the lowest natural number for which the network approached scale-free topology (*r*^*2*^ *≥* 0.9). The adjacency matrix was subsequently converted into a biologically-inspired topological overlap map (TOM), wherein pairwise gene similarities are based on comparing their connectivity profiles, a more meaningful measure than direct correlation[24]. Hierarchical clustering with superimposed hybrid adaptive tree cut[25], an unsupervised dendrogram cutting function, was applied to the TOM to identify biologically important gene modules. Finally, representative module “meta-genes” for each sample were calculated using principal component analysis; they are simply the projections of a sample’s modules’ gene expression values on their respective first principal components.

### Module investigation

We first used a multivariate logistic regression model to identify modules whose meta-gene expression levels differed significantly (p<0.05) between recurrent and non-recurrent tumors when controlling for histological grade. In order to probe the biological function of relevant modules, we used the well-established and open-source *Database for Annotation, Visualization and Integrated Discovery* (DAVID)[9] to analyze their constituent gene lists.

### Recurrence classifier

A logistic regression classifier was used as our prediction model, with module meta-genes as regressors and recurrence as a binary response. The model was trained with the discovery cohort and tested with both 10-fold cross validation and external validation sets.

### Computational platform

All analysis was performed using R, an open-source platform for statistical computation and graphics[26].

## Results

We firstly investigated the differential gene expression profile between recurrent and non-recurrent meningiomas in both cohorts, though no single genes were significantly different (log2 fold change >1 and p<0.0001) in the validation cohort (Figure 1A). Co-expression networks revealed two gene modules (Figure 1B and C) with significant correlation to recurrence (Mann-Whitney p<0.05): a “blue” module containing 220 genes and a “turquoise” module consisting of 299 genes. The meta-gene of each module was defined as the first principal component of its expression values[20](Supplementary figure 1).

**Figure 1:**
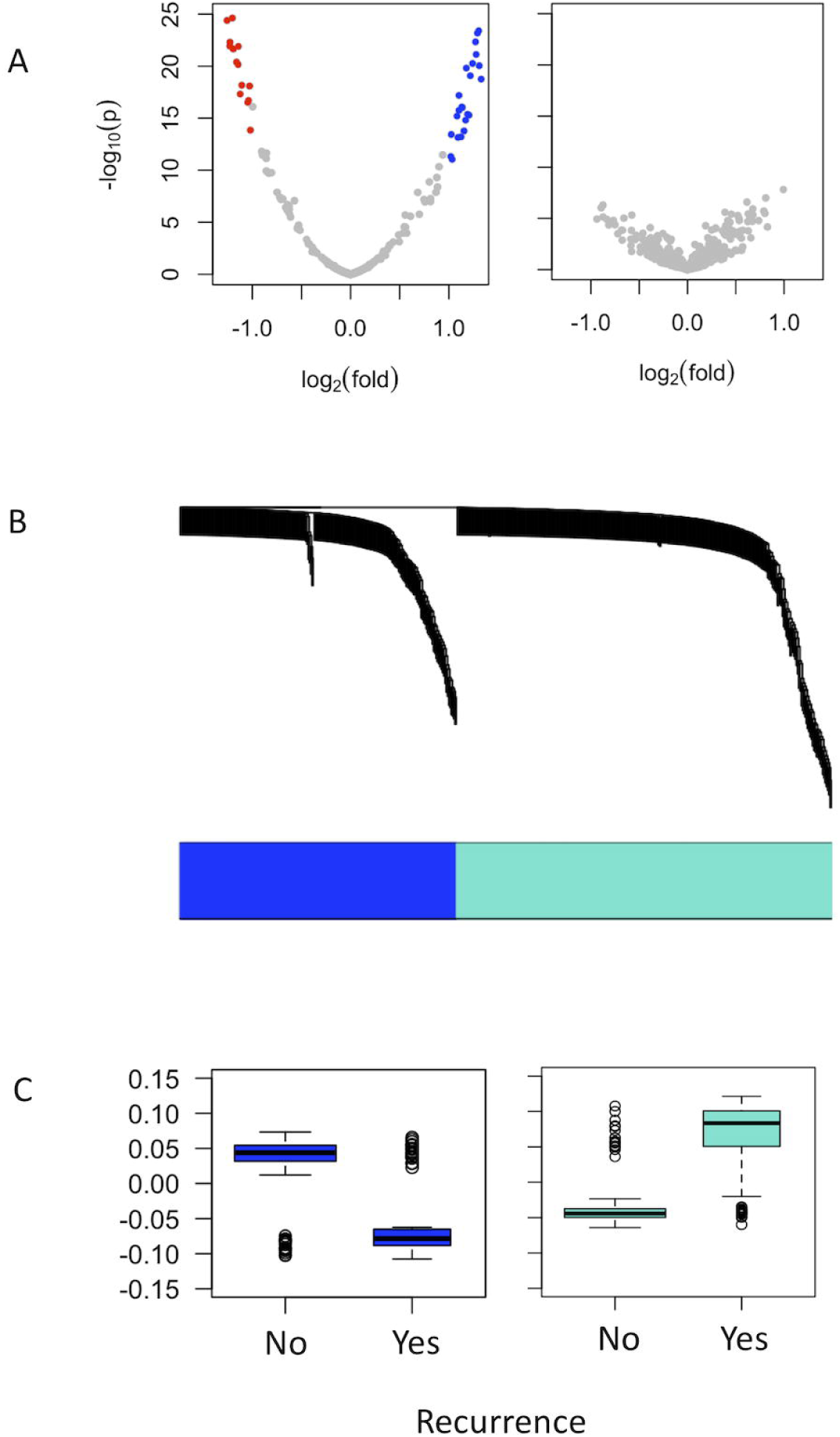
Differential gene expression and module detection. A: Differential gene expression between recurrent and non-recurrent tumors in the discovery (left) and validation (right) cohorts. Log-2 fold change and p-value cutoffs of 1, and 0.0001, respectively, were used. Significantly upregulated genes are colored blue and significantly downregulated genes are colored red. B: Dendrogram revealing 2 modules (turquoise and blue). C: Boxplots comparing meta-gene expression between recurrent and non-recurrent tumors for both modules.

We next investigated whether the meta–genes could be used to predict meningioma recurrence and found good classifier performance with 10-fold cross validation using the “turquoise” module (AUC=0.87 ±0.04; Figure 2A). Importantly, this performance was preserved in the validation cohort (AUC = 0.81 ± 0.09; Figure 2B). We also investigated the classifier performance on the subset of grade 2 tumors, since predicting recurrence in these patients is limited by particularly high uncertainty. Our classifier maintained high accuracy with 10-fold cross validation (AUC = 0.93 ± 0.05; Figure 2C), though accuracy decreased with external validation (AUC = 0.75 ± 0.22; Figure 2D), using the turquoise module. This module was enriched in biological processes related to cell division and metabolism (Figure 3A). Adding the blue module did not contribute to overall model performance. The turquoise meta-gene maintained its predictive accuracy with concurrent removal of up to 20 of the most individually predictive module genes (Figure 3C), and a meta gene built from these genes performed equally to the meta gene of the full module (Figure 3D). To ensure our model is unbiased, we randomly perturbed recurrence labels in the validation cohort, which abolished model performance as expected (Supplementary Figure 2).

**Figure 2:**
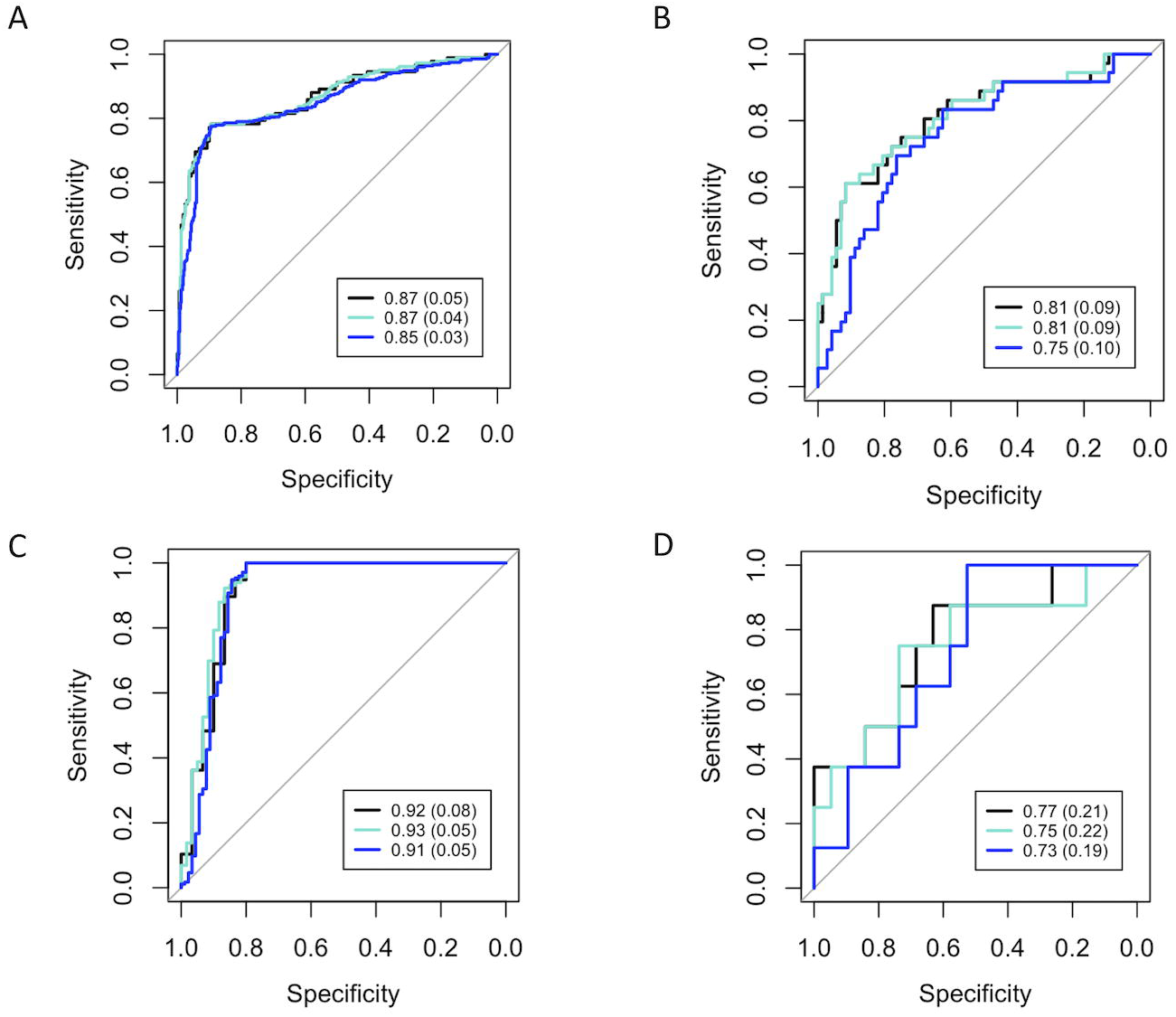
Receiver operating characteristic curves for meta-gene recurrence classifiers. Solid black curves represent a full model (both blue and turquoise meta-genes), while blue and turquoise curves represent their respective univariate models. A and B: Classifier with all grades included, using 10-fold cross validation on the discovery cohort and externally validating results on the validation cohort, respectively. C and D: Classifier ROC curves with the subset of only grade 2 tumors.

**Figure 3.**
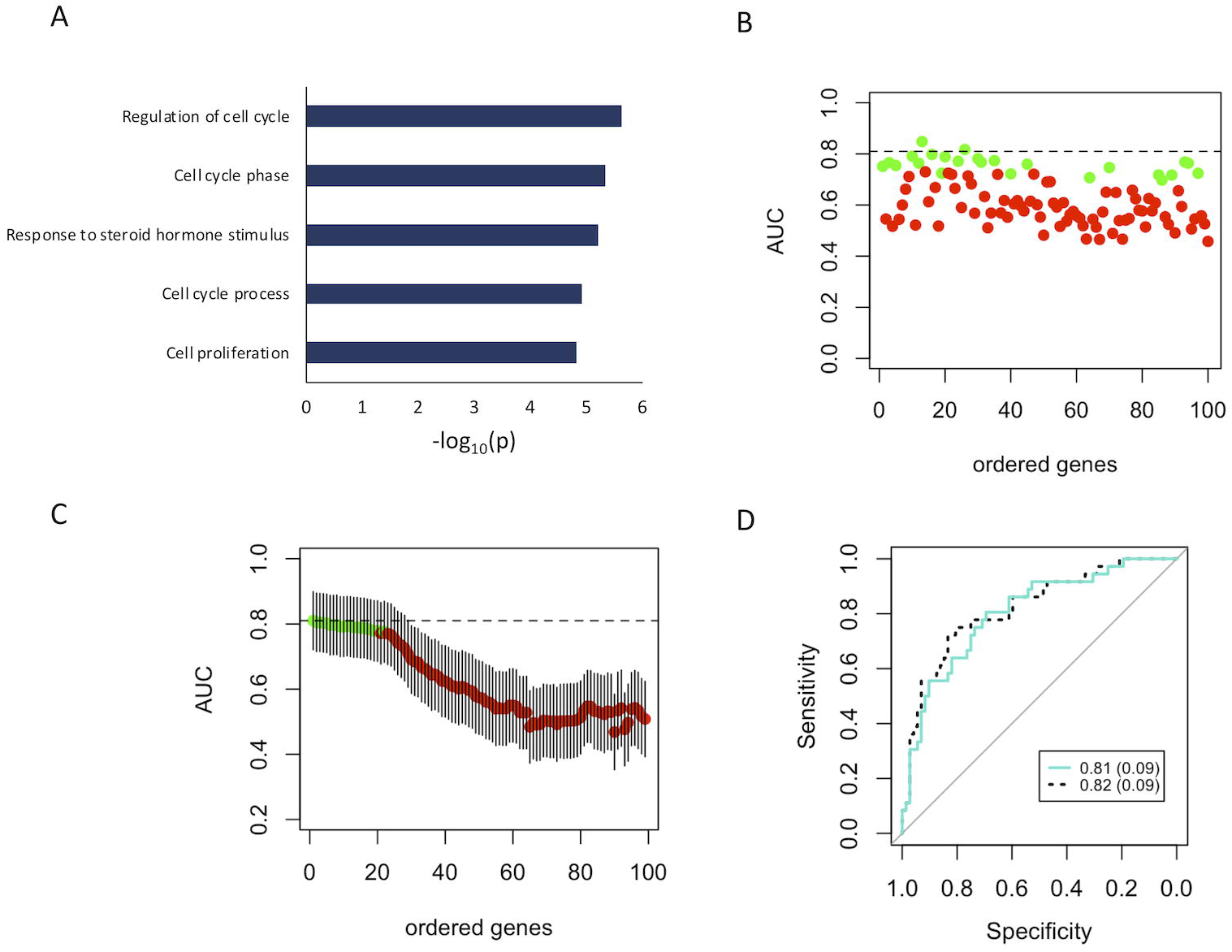
Turquoise module. A: The top 5 biological processes enriching in the turquoise module. B: Predictive accuracies of the top 100 individual genes in the module, ranked by correlation to module meta-gene. The interrupted black line represents the full module performance. Green points represent genes that whose ROC curves are not significantly different from that of the full module (DeLong’s p>0.05), while red points represent genes which are significantly different. C: AUC of ROC curves generated from iteratively removing genes in order of highest AUC, based on their single-gene validation performance (B). Twenty of the top genes must be removed before the meta-gene becomes significantly different from that of the full model. D: ROC curves comparing the full model to 20-gene sparsified model, demonstrating very similar performance.

## Discussion

We have used a simple and robust approach to predicting meningioma recurrence using gene modules, which yields high accuracy with a multi-study validation cohort. Previous analysis of the DNA methylation status of meningiomas has identified classes relevant to disease progression using unsupervised clustering. Grade 2 meningiomas were also observed to be scattered over these methylation classes further indicating diverse biology [8] and underlining the diagnostic problem created by these tumors.

In our study, we show that a systems biology approach using meta-genes provides an accurate predictor of tumor recurrence. This principle can be used in marker discovery for other challenging diseases where conventional approaches such as differential gene expression analysis have been unsuccessful. Our results are derived from data obtained under diverse conditions and yet still produce an accurate classier which projects translational value in creating a diagnostic panel.

Our study uses publicly available data and is thus limited by sparse annotation and variable follow-up times. The extent of tumor excision in our study was not known. Although this is conventionally held to affect recurrence[27] the relevance has been debated over the past decade[1] and our classifier yields high accuracy purely with gene expression data. While follow-up times are limited for some patients (Table 1) they still represent real life data from the general neurosurgical practice. Though our results are largely exploratory at present, further optimization of a predictive gene module using prospectively gathered cohorts with greater follow up times may ultimately translate into a novel approach to care of patients with meningioma at the bedside.

## Conclusions

We apply gene co-expression networks to a cohort of 252 adult meningiomas and derive a cluster of genes which is able to predict meningioma recurrence with high accuracy. With the wide accessibility of custom-made mini-arrays, our findings support, and should encourage, the shift to include this or a similar classifier in routine clinical care.

## Figure legends

**Supplementary Figure 1**: Validating the first principal component as meta-gene for predicting recurrence. A and B: Principal components two and above explain little variance of gene expression for the blue module in both training (A) and validation (B) sets. C: Graph summary of predictive accuracy (AUC) when including individual principle components as predictors in the model. D: Stepwise addition of principal component does not improve model performance.

**Supplementary Figure 2:** Classifier validity confirmed by scrambling response variable (recurrence) in validation cohort.

## Declarations

### Ethics approval and consent to participate

Not applicable, as this study relies only on pre-existing publicly available data.

### Consent for publication

Not applicable.

### Availability of data and materials

All data is available and was retrieved from the publically-accessible online genetic repository GEO omnibus (https://www.ncbi.nlm.nih.gov/geo/).

## Competing interests

The authors declare that they have no competing interests.

## Funding

Zsolt Zador: National Institute for Health Research, Academic Clinical Lectureship.

## Authors’ contributions

ZZ and AL designed the study, analyzed the data, and wrote the manuscript; BHK guided data analysis and revised the manuscript; MDC supervised the project and revised the manuscript.

## Notes

https://www.ncbi.nlm.nih.gov/geo/

## References

1. Sughrue ME, Kane AJ, Shangari G, Rutkowski MJ, McDermott MW, Berger MS, et al. The relevance of Simpson Grade I and II resection in modern neurosurgical treatment of World Health Organization Grade I meningiomas. J Neurosurg. 2010;113:1029–35.

2. Komotar RJ, Iorgulescu JB, Raper DMS, Holland EC, Beal K, Bilsky MH, et al. The role of radiotherapy following gross-total resection of atypical meningiomas. J Neurosurg. 2012;117:679–86.

3. Aghi MK, Carter BS, Cosgrove GR, Ojemann RG, Amin-Hanjani S, Martuza RL, et al. Long-term recurrence rates of atypical meningiomas after gross total resection with or without postoperative adjuvant radiation. Neurosurgery. 2009;64:56–60.

4. Vasudevan HN, Braunstein SE, Phillips JJ, Pekmezci M, Tomlin BA, Wu A, et al. Comprehensive Molecular Profiling Identifies FOXM1 as a Key Transcription Factor for Meningioma Proliferation. Cell Rep. 2018;22:3672–83.

5. Clark VE, Erson-Omay EZ, Serin A, Yin J, Cotney J, Özduman K, et al. Genomic analysis of non-NF2 meningiomas reveals mutations in TRAF7, KLF4, AKT1, and SMO. Science (80-). 2013;339:1077–80.

6. Harmancl AS, Youngblood MW, Clark VE, CoÅ Kun S, Henegariu O, Duran D, et al. Integrated genomic analyses of de novo pathways underlying atypical meningiomas. Nat Commun. 2017;8.

7. Clark VE, Harmancl AS, Bai H, Youngblood MW, Lee TI, Baranoski JF, et al. Recurrent somatic mutations in POLR2A define a distinct subset of meningiomas. Nat Genet. 2016;48:1253–9.

8. Sahm F, Schrimpf D, Stichel D, Jones DTW, Hielscher T, Schefzyk S, et al. DNA methylation-based classification and grading system for meningioma: a multicentre, retrospective analysis. Lancet Oncol. 2017;18:682–94.

9. Huang DW, Sherman BT, Lempicki RA. Systematic and integrative analysis of large gene lists using DAVID bioinformatics resources. Nat Protoc. 2009;4:44–57.

10. Vernooij M, Ikram M. Incidental findings on brain MRI in the general population. N Engl J Med. 2007;:1821–8.

11. Rogers L, Barani I, Chamberlain M, Kaley TJ, McDermott M, Raizer J, et al. Meningiomas: knowledge base, treatment outcomes, and uncertainties. A RANO review. J Neurosurg. 2015;122:4–23.

12. Jääskeläinen J. Seemingly complete removal of histologically benign intracranial meningioma: Late recurrence rate and factors predicting recurrence in 657 patients. A multivariate analysis. Surg Neurol. 1986;26:461–9.

13. Ketter R, Henn W, Niedermayer I, Steilen-Gimbel H, König J, Zang KD, et al. Predictive value of progression-associated chromosomal aberrations for the prognosis of meningiomas: a retrospective study of 198 cases. J Neurosurg. 2001;95:601–7.

14. Goutagny S, Nault JC, Mallet M, Henin D, Rossi JZ, Kalamarides M. High incidence of activating TERT promoter mutations in meningiomas undergoing malignant progression. Brain Pathol. 2014;24:184–9.

15. Yesilöz Ü, Kirches E, Hartmann C, Scholz J, Kropf S, Sahm F, et al. Frequent AKT1E17Kmutations in skull base meningiomas are associated with mTOR and ERK1/2 activation and reduced time to tumor recurrence. Neuro Oncol. 2017;19:1088–96.

16. Boetto J, Bielle F, Sanson M, Peyre M, Kalamarides M. SMO mutation status defines a distinct and frequent Molecular subgroup in olfactory groove meningiomas. Neuro Oncol. 2017;19:345–51.

17. Ghazalpour A, Doss S, Zhang B, Wang S, Plaisier C, Castellanos R, et al. Integrating genetic and network analysis to characterize genes related to mouse weight. PLoS Genet. 2006;2:1182–92.

18. Horvath S, Zhang B, Carlson M, Lu K V., Zhu S, Felciano RM, et al. Analysis of oncogenic signaling networks in glioblastoma identifies ASPM as a molecular target. Proc Natl Acad Sci. 2006;103:17402–7.

19. Langfelder P, Cantle JP, Chatzopoulou D, Wang N, Gao F, Al-Ramahi I, et al. Integrated genomics and proteomics define huntingtin CAG length-dependent networks in mice. Nat Neurosci. 2016;19:623–33.

20. Langfelder P, Horvath S. WGCNA: An R package for weighted correlation network analysis. BMC Bioinformatics. 2008;9.

21. Barrett T, Wilhite SE, Ledoux P, Evangelista C, Kim IF, Tomashevsky M, et al. NCBI GEO⍰: archive for functional genomics data sets — update. Nucleic Acids Res. 2013;41 June:991–5.

22. Chen C, Grennan K, Badner J, Zhang D, Gershon E, Jin L, et al. Removing Batch Effects in Analysis of Expression Microarray Data⍰: An Evaluation of Six Batch Adjustment Methods. PLoS One. 2011;6.

23. Hughey JJ, Butte AJ. Robust meta-analysis of gene expression using the elastic net. Nucleic Acids Res. 2015;43:1–11.

24. Zhang B, Horvath S. A General Framework for Weighted Gene Co-expression Network Analysis. Stat Appl Genet Mol Biol. 2005;4.

25. Methods T, Clustering H, Author D, Langfelder P, Zhang B, Horvath S, et al. Package ‘dynamicTreeCut.’ 2016;:1–14.

26. R Development Core Team. R: A Language and Environment for Statistical Computing. R Found Stat Comput Vienna Austria. 2016;0:{ISBN} 3-900051-07-0. doi:10.1038/sj.hdy.6800737.

27. Simpson D. The recurrence of intracranlal meningiomas after surgical treatment. J Neurol Neurosurg Psychiatry. 1957;20:22–39.

