## Supplementary figures and images for "Meta-gene markers predict meningioma recurrence with high accuracy"

### Supplemental figures

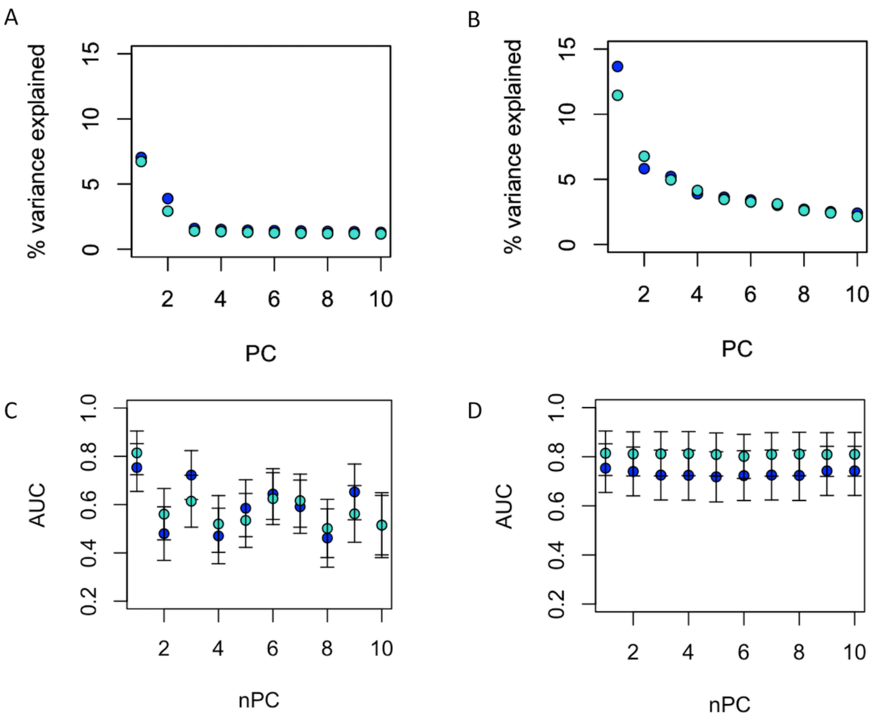

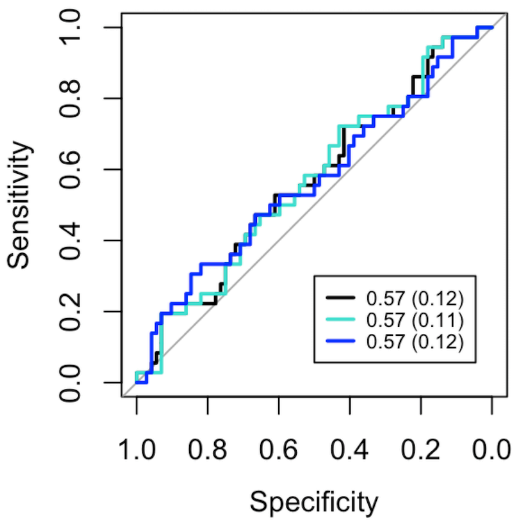
